# Loss of function variants in *PCYT1A* causing spondylometaphyseal dysplasia with cone/rod dystrophy have broad consequences on lipid metabolism, chondrocyte differentiation, and lipid droplet formation

**DOI:** 10.1101/2019.12.19.882191

**Authors:** Julie Jurgens, Suming Chen, Nara Sobreira, Sarah Robbins, Arianna Franca Anzmann, Raha Dastgheyb, Saja S. Khuder, Julie Hoover-Fong, Courtney Woods, Felicity Collins, John Christodoulou, Guilherme Lopes Yamamoto, Débora Romeo Bertola, Wagner A. R. Baratela, Sophie D. Curie, Norman Haughey, Rosemary Cornell, David Valle

## Abstract

**Abstract:** Spondylometaphyseal dysplasia with cone-rod dystrophy (SMD-CRD) is a rare autosomal recessive disorder of the skeleton and the retina caused by biallelic variants in *PCYT1A*, encoding the nuclear enzyme CTP:phosphocholine cytidylyltransferase α (CCTα), which catalyzes the rate-limiting step in phosphatidylcholine (PC) biosynthesis by the Kennedy pathway. As a first step in understanding the consequences of *PCYT1A* variants on SMD-CRD pathophysiology, we generated and characterized a series of cellular models for SMD-CRD, including CRISPR-edited *PCYT1A*-null HEK293 and ATDC5 cell lines. Immunoblot and PC synthesis assays of cultured skin fibroblasts from SMD-CRD patient cell lines revealed patient genotype-specific reductions in CCTα steady state levels (10-75% of wild-type) and choline incorporation into PC (22-54% of wild-type). While *PCYT1A*-null HEK293 cells exhibited fewer and larger lipid droplets in response to oleate loading than their wild-type counterparts, SMD-CRD patient fibroblasts (p.Ser323Argfs*38 homozygotes) failed to show significant differences in lipid droplet numbers or sizes as compared to controls. Lipid droplet phenotypes in *PCYT1A*-null HEK293 cells were rescued by transfection with wild-type, p.Ala99Val, and p.Tyr240His human *PCYT1A* cDNAs. While both edited cellular models had normal morphology and proliferation rates compared to unedited controls, *Pcyt1a*-null ATDC5 cells demonstrated accelerated rates of chondrocyte differentiation as compared to their wild-type counterparts. Lipidomics revealed changes in 75-200 lipid levels in *PCYT1A*-null HEK293 and ATDC5 cells or in SMD-CRD patient fibroblasts as compared to wild-type controls. The specific lipids altered and extent of change varied by cell type. Importantly, both *PCYT1A*-null HEK293 cells and SMD-CRD patient fibroblast cell lines had decreased phosphatidylcholine:phosphatidylethanolamine (PC:PE) ratios and decreased levels of several lysophosphatidylcholine (LPC) species as compared to wild-type controls, suggesting compensatory PC production through increased LPC remodeling by LPCAT or decreased conversion of PC to LPC by phospholipase A_2_. Our results show that all tested *PCYT1A* alleles associated with SMD-CRD are hypomorphic and suggest involvement of *PCYT1A* in chondrocyte differentiation, PC:PE ratio maintenance and LPC metabolism, and lipid droplet formation.

**Author Summary:** Rare genetic disorders can reveal the function of genes on an organismal scale. When normal gene activity is lost, patients can experience a range of symptoms, often dependent on the residual activity of the encoded protein. Rare variants in the gene *PCYT1A* can cause multiple inherited disorders, including a disorder of the skeleton and the retina characterized by short stature, bone abnormalities, and blindness. *PCYT1A* is required for normal cellular function, particularly lipid metabolism, but the role of this gene in human disease is still poorly understood. To determine consequences of genetic variants in patients with this disorder, we made and studied a series of cellular models, including cells cultured from patients and CRISPR-edited cell lines lacking normal copies of *PCYT1A*. Here we show that patient variants lead to reduced *PCYT1A* expression and/or function and have adverse consequences on cell biology and lipid metabolism that are often cell-type specific. This work advances understanding of the role of lipid metabolism in skeletal and eye development.

## Introduction

Spondylometaphyseal dysplasia with cone-rod dystrophy (SMD-CRD, MIM 608940) is a rare, autosomal recessive disorder of the skeleton and the retina. The clinical phenotype includes progressive, early-onset photoreceptor degeneration—particularly in the macula—as well as short stature, bowing of the long bones, metaphyseal flaring, rhizomelic shortening, platyspondyly, and scoliosis. Using whole exome and Sanger sequencing, Hoover-Fong et al. (2014) and Yamamoto et al. (2014) identified homozygous or compound heterozygous variants in *PCYT1A* as the cause of SMD-CRD [1,2]. These variants ranged from missense (RefSeq NM_005017.2: c.296C>T, p.Ala99Val; c.295G>A, p.Ala99Thr; c.385G>A, p.Glu129Lys; c.448C>G, p.Pro150Ala; c.571T>C, p.Phe191Leu; and c.669G>C, p.Arg223Ser) to nonsense variants (c.847C>T, p.Arg283*) or frameshifting indels (c.990delC, p.Ser331Profs*?; c.968dupG, p.Ser323Argfs*38; c.996delC, p.Ser333Leufs*?). Since the publication of these initial reports, our group has identified two additional missense alleles in *PCYT1A* in patients with SMD-CRD (c.341G>C, p.Ser114Thr and c.718T>C, p.Tyr240His).

Interestingly, compound heterozygous *PCYT1A* variants were also detected in two unrelated young adult probands with congenital lipodystrophy and fatty liver disease [3]. These patients had no detectable retinal or skeletal abnormalities, suggesting a distinct pathophysiological mechanism. Both patients shared an in-frame deletion variant (c.838_840delCTC, p.Glu280del) in trans to a second variant in the other allele (c.424G>A, p.Val142Met or c.996delC, p.Ser333Leufs*?, respectively). These observations suggest that the shared allele may account for the distinctive phenotype in these patients. Testa et al. (2017) reported a third class of patients with biallelic *PCYT1A* variants and retinal degeneration, without accompanying skeletal dysplasia or lipodystrophy [4]. The two probands shared a common *PCYT1A* missense variant (c.277G>A, p.Ala93Thr) in trans to a splice site or nonsense variant in the second allele (c.897+1G>A or c.847C>T, p.Arg283*, respectively).

*PCYT1A* encodes CTP:phosphocholine cytidylyltransferase α (CCTα), which catalyzes the rate-limiting step in *de novo* synthesis of phosphatidylcholine (PC) by the Kennedy pathway [5]. *PCYT1A* is expressed ubiquitously in humans and mice, and PC is the predominant membrane phospholipid in mammalian cells [6–10]. Complete loss of *Pcyt1a* expression has devastating consequences: homozygosity for a *Pcyt1a* knockout allele results in embryonic lethality by day E3.5 and failed implantation [9], and MT-58 CHO cells with a temperature-sensitive *Pcyt1a* point mutation resulting in undetectable CCT protein at 40°C undergo apoptosis at this temperature [11,12]. Based on this evidence, some degree of PC production by the Kennedy pathway appears to be essential for life, at least in these model systems. Fly models with eye-specific RNAi knockdown or ocular mosaic homozygous knockout of *CCT1* (the fly ortholog of *PCYT1A*) showed reduced response to light on ERG, decreased rhodopsin expression, and failed rhabdomere formation [13]. These phenotypes were rescued with wild-type and nuclear localization signal mutant *CCT1*. In at least some cells, however, *PCYT1A* is expressed at levels exceeding that required for basal function. For example, MT-58 CHO cells at the permissive temperature have normal growth rates with only ~5% CCT expression compared to control CHO cells [12].

Although most PC is produced by the CCTα-dependent Kennedy pathway in mammalian cells, there are alternative pathways for PC production [6,14] (Fig 1). *PCYT1B*, a paralog of *PCYT1A*, encodes the enzyme CCTβ, which catalyzes the same Kennedy pathway reaction as CCTα but has expression limited predominantly to the brain and reproductive tissues of mice and humans [7,10,14]. Independent of the Kennedy pathway, PC can be formed by sequential methylation of another phospholipid, phosphatidylethanolamine (PE), by the PEMT pathway, which functions primarily in liver and adipocytes [5,15]. Finally, in a series of reactions known as the Lands cycle, PC can be converted into lysophosphatidylcholine (LPC) through fatty acid hydrolysis catalyzed by phospholipase A_2_ (PLA_2_), and LPC can be re-acylated to form PC by lysophosphatidylcholine acyltransferases (LPCATs) [16,17]. Interestingly, rd11 mice with *Lpcat1* deletions have a retinal degeneration phenotype, bolstering support for involvement of this arm of the phospholipid metabolism pathway in retinal degeneration [18]. Perturbations in other Kennedy pathway genes have been linked to phenotypes independent of the skeleton or retina, suggesting a versatile role for lipid metabolism [19–21].

**Fig 1.**
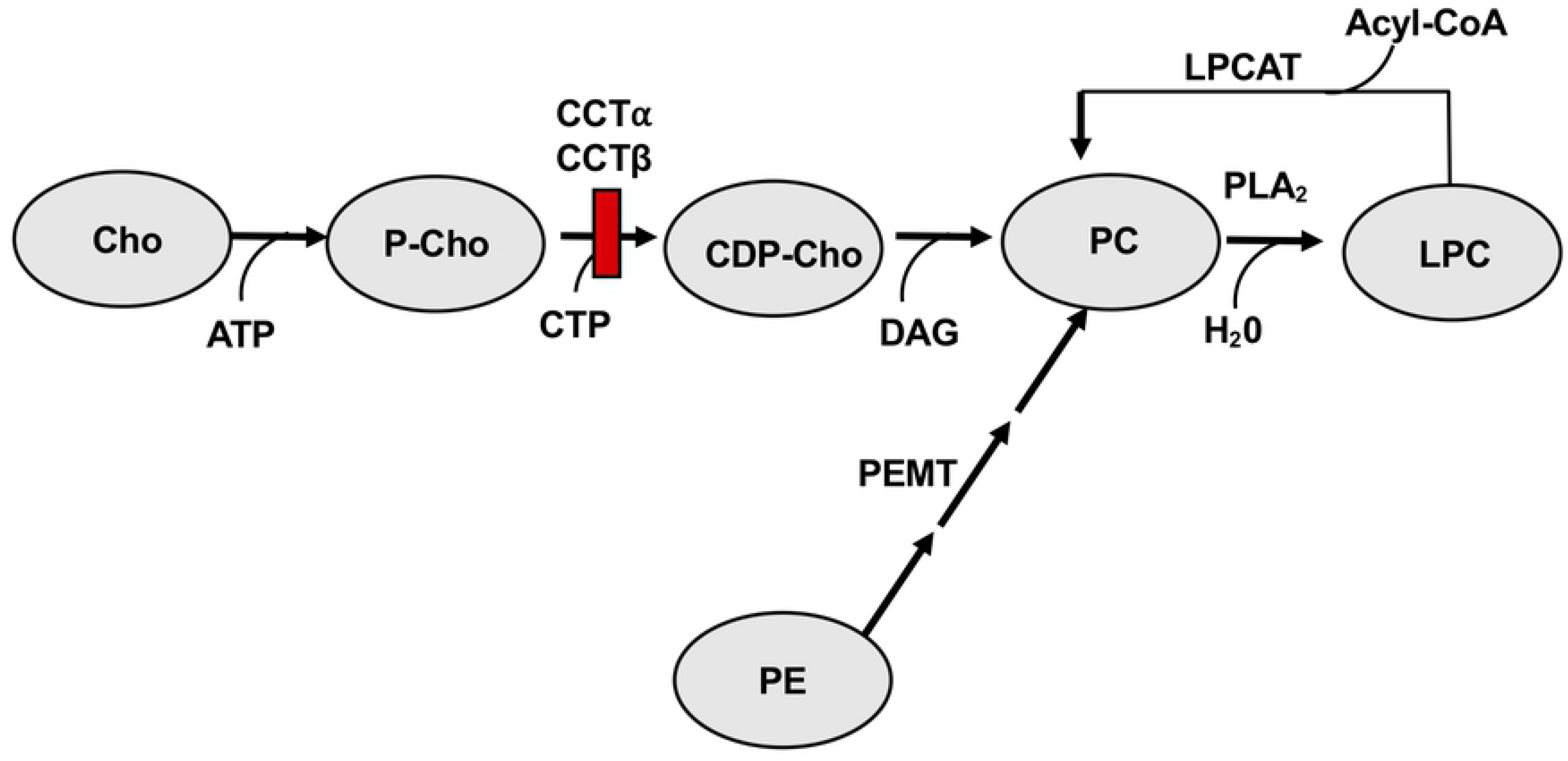
Kennedy pathway and related pathways for PC production. Choline (Cho) is taken in exogenously from the diet and transported across the plasma membrane into the cytosol, where it becomes phosphorylated by choline kinase to produce phosphocholine (P-Cho). Phosphocholine is converted into CDP-choline (CDP-Cho) in a reaction catalyzed by CTP:phosphocholine cytidylyltransferase (CCT), which is combined with diacylglycerol (DAG) to form phosphatidylcholine (PC). Phospholipase-A_2_ (PLA_2_) converts PC into lyso-phosphatidylcholine (LPC), which can be converted back into PC in a reaction catalyzed by lysophosphatidylcholine acyltransferase (LPCAT). PC can also be derived from sequential methylation of phosphatidylethanolamine (PE) by phosphatidylethanolamine *N*-methyltransferase. The red rectangle indicates the location of the block in PC synthesis in SMD-CRD. Phosphatidylethanolamine can also be interconverted to lysophosphatidylethanolamine by phospholipase-A_2_ (PLA_2_) and LPE acyltransferase (LPEAT) enzymes.

CCTα functions as a homodimer comprised of 367-residue monomers with several functional domains (Fig 1) [6]. The catalytic domain is responsible for catalyzing the conversion of phosphocholine and CTP to CDP-choline and diphosphate. In addition, there is an N-terminal nuclear localization signal; a membrane-binding amphipathic helical domain (M domain); and a C-terminal phosphorylation domain, which when phosphorylated reduces membrane association *in vitro* and in cultured cells [22]. Membrane binding of domain M in response to altered membrane lipid composition elicits catalytic activation of the enzyme by displacing an auto-inhibitory segment of the M domain from the base of the active site [6]. Immunostaining experiments have shown that CCTα in the nucleoplasm translocates to the inner nuclear membrane in response to demand for lipid synthesis or remodeling, or in response to requirements for lipid droplet expansion in yeast, fly, and in multiple types of mammalian cells, including adult mouse photoreceptors, hypertrophic zone chondrocytes from developing mouse growth plates, hepatocytes, and adipocytes [13,23,24]. Most SMD-CRD variants are missense mutations in the catalytic domain of the enzyme (Fig 2), although two variants are in the M domain (p.Tyr240His and p.Arg283*) and three frameshifting indels localize to the C-terminal phosphorylation domain (p.Ser331Profs*?, p.Ser323Argfs*38, and p.Ser333Leufs*?). The catalytic domain mutant enzymes have defects in folding stability based on thermal denaturation of purified enzymes, and several have impaired catalytic function *in vitro*. For example, p.Ala99Val, p.Ala99Thr, p.Glu129Lys, and p.Arg223Ser have higher K_m_ values for both substrates, CTP and phosphocholine, and lower V_max_ values, suggesting that in cells expressing these enzymes the rate of CDP-choline production would be slower [25].

**Fig 2.**
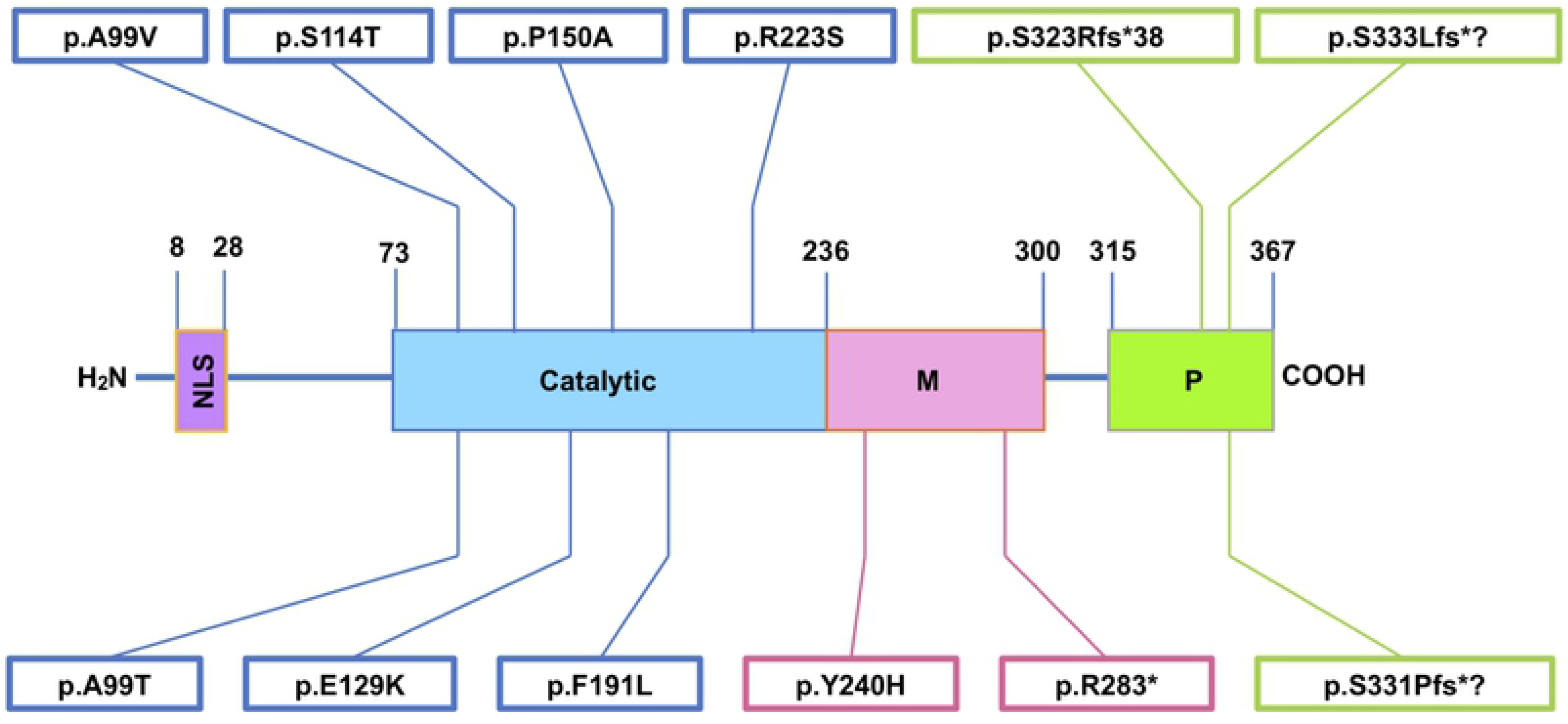
Diagram of all known SMD-CRD alleles to-date. Both previously described and novel SMD-CRD patient variants are mapped to their corresponding CCTα protein domains. Note that most variants are missense and fall within the catalytic domain. NLS-nuclear localization signal; M-membrane-binding domain; P-phosphorylation domain.

An additional aspect of CCTα cell biology is its relationship with cellular organelles called lipid droplets (LDs). LDs form in response to loading of cells with fatty acids (e.g. oleate) that are converted to triglycerides for storage, and to prevent lipotoxicity [26]. Phospholipids such as PC are required for the formation of LD membranes, which encapsulate neutral lipids and prevent their coalescence [27]. Neutral lipids such as fatty acids and diacylglycerol activate CCTα and stimulate PC production; for instance, oleate facilitates the dephosphorylation and membrane translocation of CCTα, leading to its activation [23,24,28–31]. Accordingly, CCTα-deficient yeast, *Drosophila*, mouse, rat, and human cells accumulate fewer and larger lipid droplets as compared to their wild-type counterparts in response to oleate loading [13,24,27,32,33].

Here we describe the generation and characterization of three distinct cellular model systems—cultured human dermal fibroblasts, HEK293 cells, and mouse pre-chondrocyte ATDC5 cells—to interrogate diverse functional consequences of *PCYT1A* loss of function. Using cultured skin fibroblasts obtained from SMD-CRD patients, we sought to understand the consequences of SMD-CRD variants on cellular phenotypes in an endogenous context. Next, we generated *PCYT1A*-null HEK293 cell lines to assess the impact of complete loss of CCTα on previously described CCT functions such as lipid droplet formation. Finally, we created *Pcyt1a*-null ATDC5 cell lines, which are putatively more relevant to SMD-CRD pathophysiology, to measure the effects of loss of CCTα on chondrocyte proliferation and differentiation.

## Results

### Measurement of CCTα steady state levels and PC synthesis rates

Immunoblot analysis of CCTα steady state levels in cultured fibroblasts of five SMD-CRD probands and one affected sibling revealed reduced CCTα steady state levels as compared to wild-type controls, with the level of reduction varying according to patient genotype (Fig 3A). Fibroblasts homozygous for the p.Ala99Val variant had the mildest reductions in CCTα steady state levels (~75% wild-type levels), followed by p.Glu129Lys homozygotes (~30% wild-type levels). Fibroblasts homozygous for a 1-bp frameshifting insertion (p.Ser323Argfs*38) had only ~10% of wild-type CCTα levels, and similar levels were observed in cells compound heterozygous for the p.Ser323Argfs*38/p.Ser114Thr variants.

**Fig 3.**
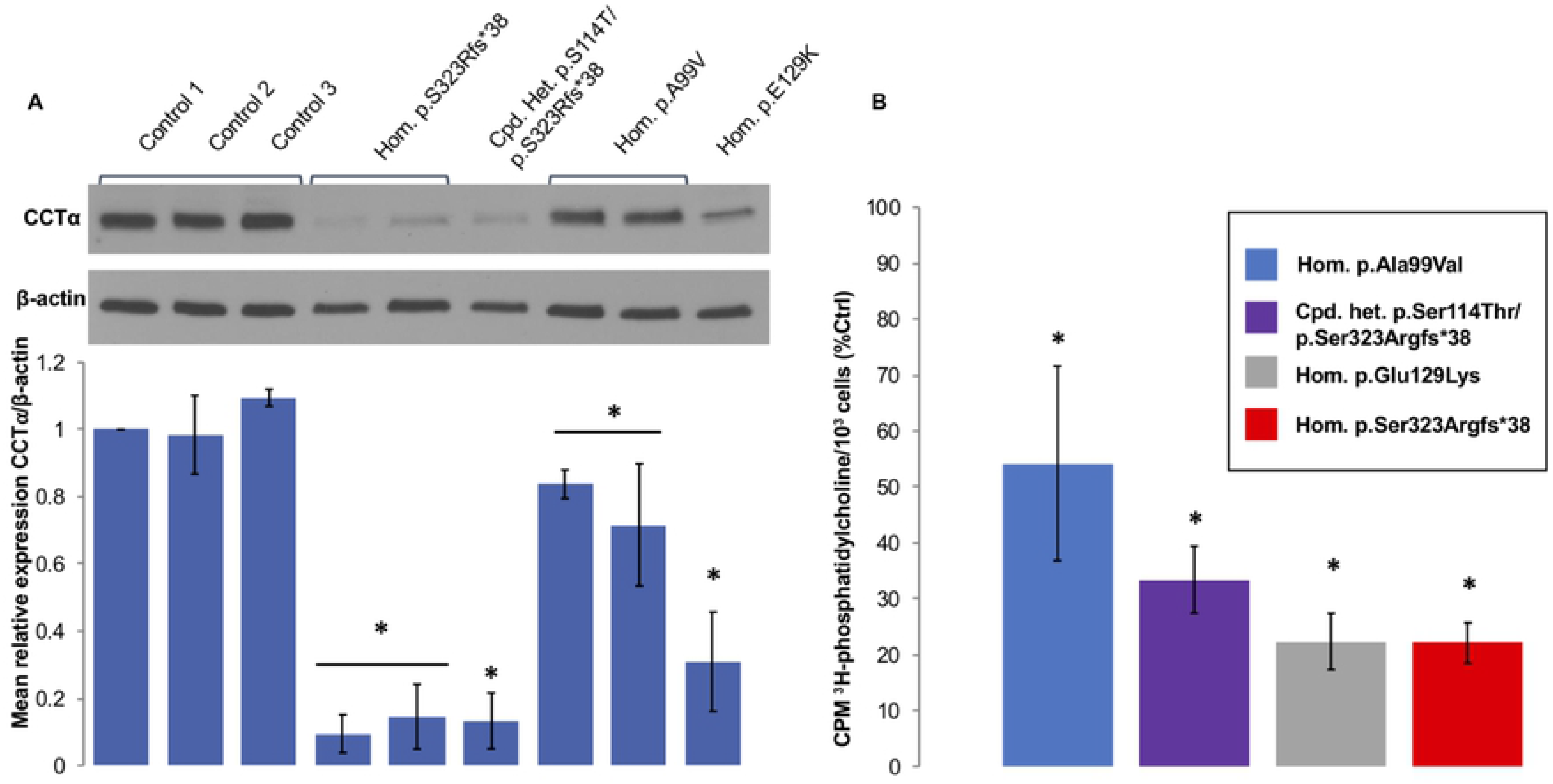
Western blot and PC synthesis assays of SMD-CRD patient fibroblasts. A) Representative immunoblot showing reduced protein expression of CCTα relative to loading control β-actin in cultured skin fibroblasts of SMD-CRD patients as compared to wild-type controls, with the level of reduction varying according to patient genotype. B) To assess PC synthesis, we measured continuous incorporation of [^3^H]-choline into PC in intact fibroblasts over 2 hours. Error bars represent standard deviation. n=3 biological replicates. Tukey’s Honest Significance Test. *p<0.05.

To assess PC synthesis, we measured continuous incorporation of [^3^H]-choline into PC in intact fibroblasts over 2 hours. Consistent with the results of our western blotting experiments, p.Ala99Val homozygotes had the mildest impairments in PC synthesis (54% of wild-type), whereas homozygotes for the p.Ser323Argfs*38 or p.Glu129Lys alleles had only 22% of wild-type incorporation levels, and compound heterozygotes for the p.Ser323Argfs*38/p.Ser114Thr alleles had 33% of wild-type incorporation.

### CCTα membrane translocation in putative cell models

In order to assess the utility of several cell models for studying CCTα variants, we first analyzed CCTα localization in response to oleate treatment. Previous studies have shown that oleate stimulation can induce CCTα translocation to the inner nuclear membrane, colocalizing with lamin A/C in CHO cells and in HEK293 cells, although results are variable depending on the report and the cell line tested [23,24,29]. Under our experimental conditions, oleate failed to stimulate translocation of CCTα to nuclear membranes in wild-type fibroblasts, but did induce nuclear membrane translocation in wild-type HEK293 cells (S1 Fig). These results are consistent with the observations of Aitchison et al., who demonstrated oleate-dependent translocation of CCTα to the nuclear membrane in HEK293 cells but not in CHO cells [24]. Based on this finding, we performed additional cellular modeling in HEK293 cells to examine consequences of *PCYT1A* variants on cellular phenotypes.

### Lipid droplet analyses

In multiple species, cellular deficiency of CCTα orthologs results in formation of fewer and larger lipid droplets than in control cells in response to oleate exposure [24,27,32,33]. We hypothesized that SMD-CRD patient fibroblast cells would have a similar phenotype in response to oleate treatment. However, we found the sizes and numbers of lipid droplets in both wild-type and SMD-CRD patient fibroblast cell lines were highly heterogeneous, and SMD-CRD patient fibroblasts (p.Ser323Argfs*38 homozygotes) failed to show significant differences in lipid droplet numbers or sizes as compared to control cells (Fig 4A and 4C). To assess the impact of total *PCYT1A* deficiency we generated *PCYT1A*-null HEK293 cells (S2 Fig). Interestingly, these cells generated fewer and larger lipid droplets than their wild-type counterparts in response to oleate loading (Fig 4B and 4D). Transfection with wild-type, p.Ala99Val, and p.Tyr240His *PCYT1A* cDNAs rescued lipid droplet phenotypes in *PCYT1A*-null cells (Fig 4E). This result suggests that either alleles with partial CCTα activity are sufficient to support normal lipid droplet morphology in response to oleate loading in HEK293 cells, or that overexpression of these hypomorphic alleles is sufficient for phenotypic rescue.

**Fig 4.**
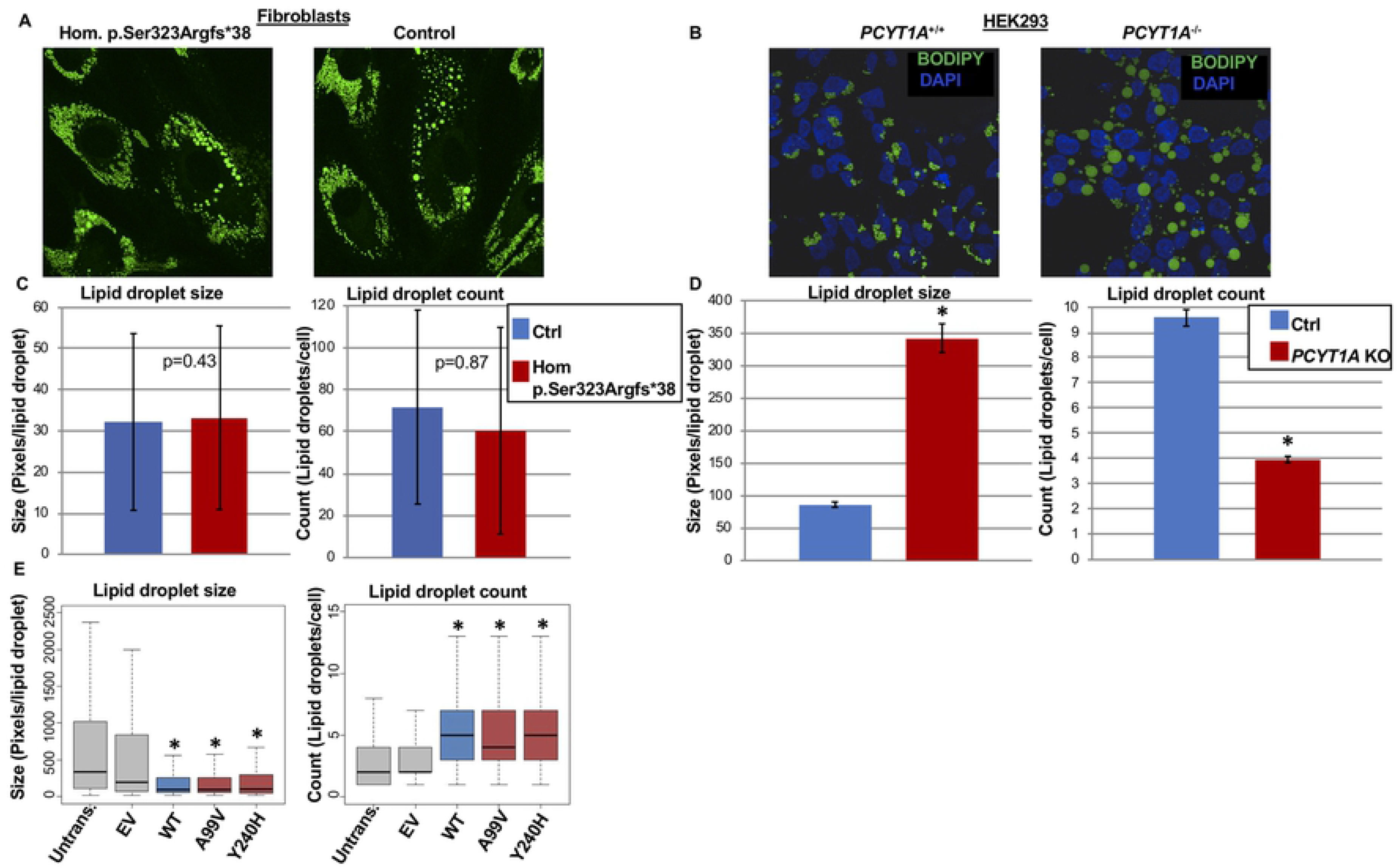
Lipid droplet phenotype and rescue experiments. Lipid droplet (LD) formation was induced by oleic acid loading (1 mM, 24 hr) and visualized with BODIPY staining. LD size and number were quantified using “analyze particle” command in ImageJ (v1.47). Note heterogeneity in LDs. A and C) Neither LD size (p=0.43) nor number (p=0.87) differed between control fibroblasts and SMD-CRD patient fibroblasts. B and D) *PCYT1A*-null HEK293 cells have fewer and larger LDs than control HEK293 cells. E) LD sizes and numbers are rescued by transfection of *PCYT1A*-null HEK293T cells with *PCYT1A* WT, p.Ala99Val or p.Tyr240His cDNAs but not corresponding empty vector controls. n=200 HEK293 *PCYT1A* null cells from 4 experimental trials. Error bars represent SD. We used Welch Two Sample t-test for panels C and D, and Tukey’s Honest Significant Difference test for panel E. Data in panel E is represented as box and whisker plot for which the box represents the range of the first to third quartiles, line represents the median, and whiskers extend to minimum and maximum data points. *p<0.05.

### Proliferation studies in HEK293 cells

Given the high concentration of PC in plasma membranes of cultured cells, we hypothesized that CCTα deficiency would limit their proliferation. Interestingly, we found that *PCYT1A*-null HEK293 cells proliferated at normal rates as compared to wild-type cells cultured in standard medium supplemented with 10% FBS (Fig 5A). To determine if this normal growth rate was dependent on uptake of exogenous lipid from the medium, we also tested cells cultured in medium supplemented with 10% delipidated serum. Both wild-type and *PCYT1A*-null HEK293 cells grew more slowly in medium with delipidated serum as compared to normal serum, but surprisingly the *PCYT1A*-null HEK293 cells grew at a faster rate than their wild-type counterparts in medium supplemented with delipidated serum and reached numbers comparable to cells grown in standard serum by day 6 (Fig 5A).

**Fig 5.**
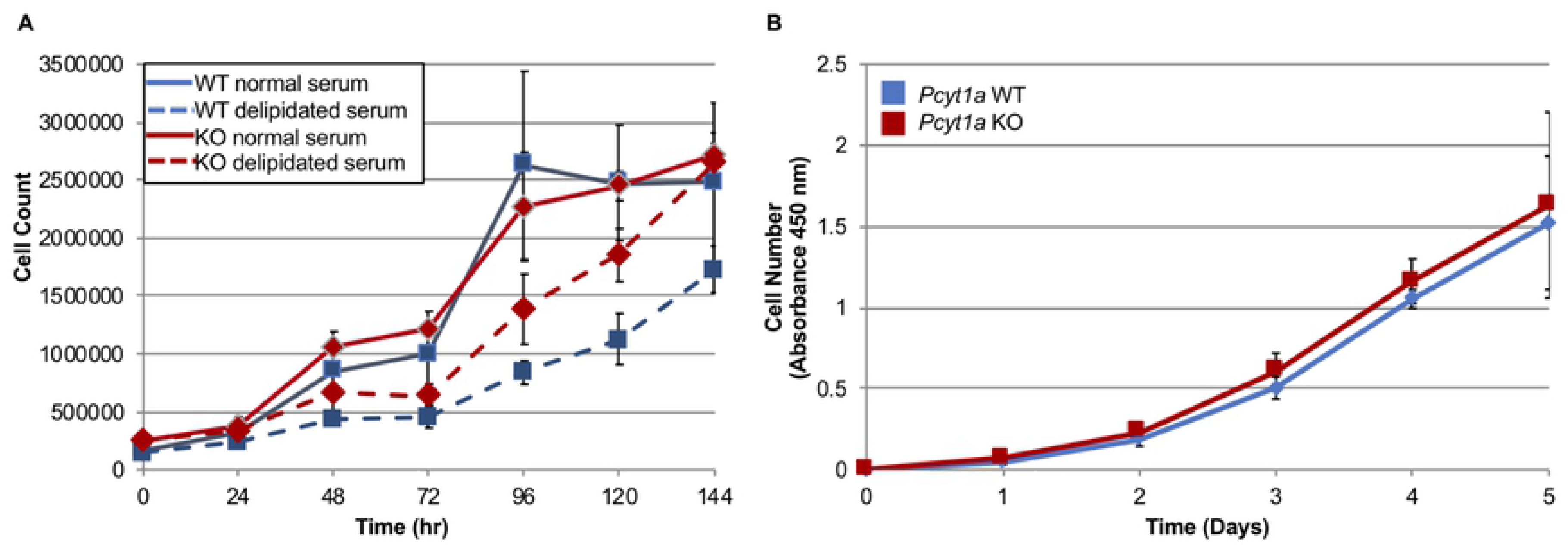
*PCYT1A*-null HEK293 and ATDC5 cells have normal proliferation rates. A) Wild-type and *PCYT1A*-null HEK293 cells were grown in either normal serum or delipidated serum up to 144 hours and counted every 24 hours. Error bars represent SD. B) Undifferentiated wild-type or *Pcyt1a*-null ATDC5 cells were grown for 5 days in normal serum and counted every 24 hours using absorbance-based cell counting. Error bars represent SD of three replicates.

### Effects of *Pcyt1a* knockout on chondrocyte proliferation and differentiation

To better understand the chondrocyte defects in SMD-CRD, we evaluated the utility of ATDC5 pre-chondrocytes as a model system. We first assessed steady-state CCTα levels at baseline and over the course of 21 days of chondrocyte differentiation by immunoblot. CCTα protein was expressed robustly in ATDC5 cells prior to chondrocyte differentiation and maintained stable expression over the 21-day differentiation time course (Fig 6A). Next, we generated *Pcyt1a*-null ATDC5 cell lines using genome editing (S2 Fig) and assessed the consequences of *Pcyt1a* loss-of-function on chondrocyte proliferation and differentiation. As in HEK293 cells, we found *Pcyt1a*-null ATDC5 cells proliferated at normal rates as compared to their wild-type counterparts (Fig 5B). Surprisingly, *Pcyt1a*-null ATDC5 cells differentiated into chondrocytes more quickly than their wild-type counterparts (Fig 6B).

**Fig 6.**
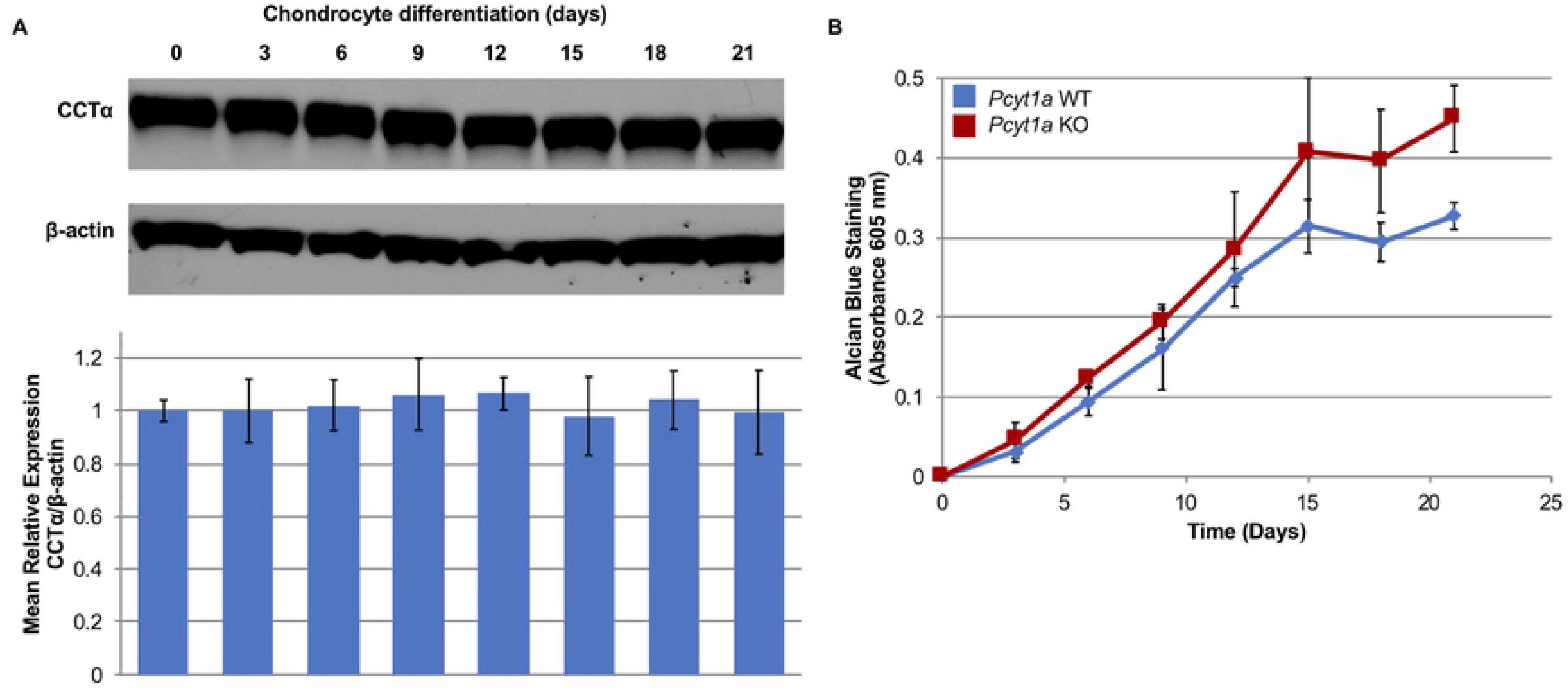
Involvement of Pcyt1a in ATDC5 chondrocyte differentiation. A) Western blot analysis of ATDC5 cell lysates shows that CCTα steady-state levels remain stable over the course of 21 days of chondrocyte differentiation. B) Rates of chondrocyte differentiation in control and *Pcyt1a*-null ATDC5 cells were measured by absorbance of Alcian blue in SDS-lysed cell extracts. n=3 biological replicates for each time point.

### Lipidomic studies

To assess the broader consequences of CCTα deficiency on cellular lipid metabolism, we performed lipidomics on ~1000 distinct lipids extracted from SMD-CRD patient-derived skin fibroblasts (p.Ser323Argfs*38 homozygotes) and in *PCYT1A*-null HEK293 or ATDC5 cells as compared to their wild-type counterparts. The SMD-CRD genotype we chose had the strongest impact on CCTα expression and PC synthesis, so we expected a stronger effect on lipid composition. We grouped lipid species distinguished by acyl/alkyl chain content into the following classes for analysis: cholesterol ester, ceramide, desmosteryl ester, diacylglycerol, galactosylceramide, lactosylceramide, lysophosphatidylcholine, lysophosphatidylethanolamine, lysophosphatidylglycerol, lysophosphatidylserine, monoakyldiacylglyceride, phosphatidic acid, phosphatidylcholine, phosphatidylethanolamine, phosphatidylglycerol, phosphatidylserine, sphingomyelin, or triacylglyceride. Some lipid classes were not represented in all cell lines tested. In total, the levels of ~75-200 lipids were significantly changed in the *PCYT1A*-null or SMD-CRD cells, and the specific lipids altered were largely variable depending on the cell type tested (S2 Table, S3 Table, S4 Table). However, there were two significant changes in the lipidome shared by both SMD-CRD cells and *PCYT1A*-null cells: reduced LPC content and reduced phosphatidylcholine: phosphatidylethanolamine (PC:PE) ratios (Fig 7).

**Fig 7.**
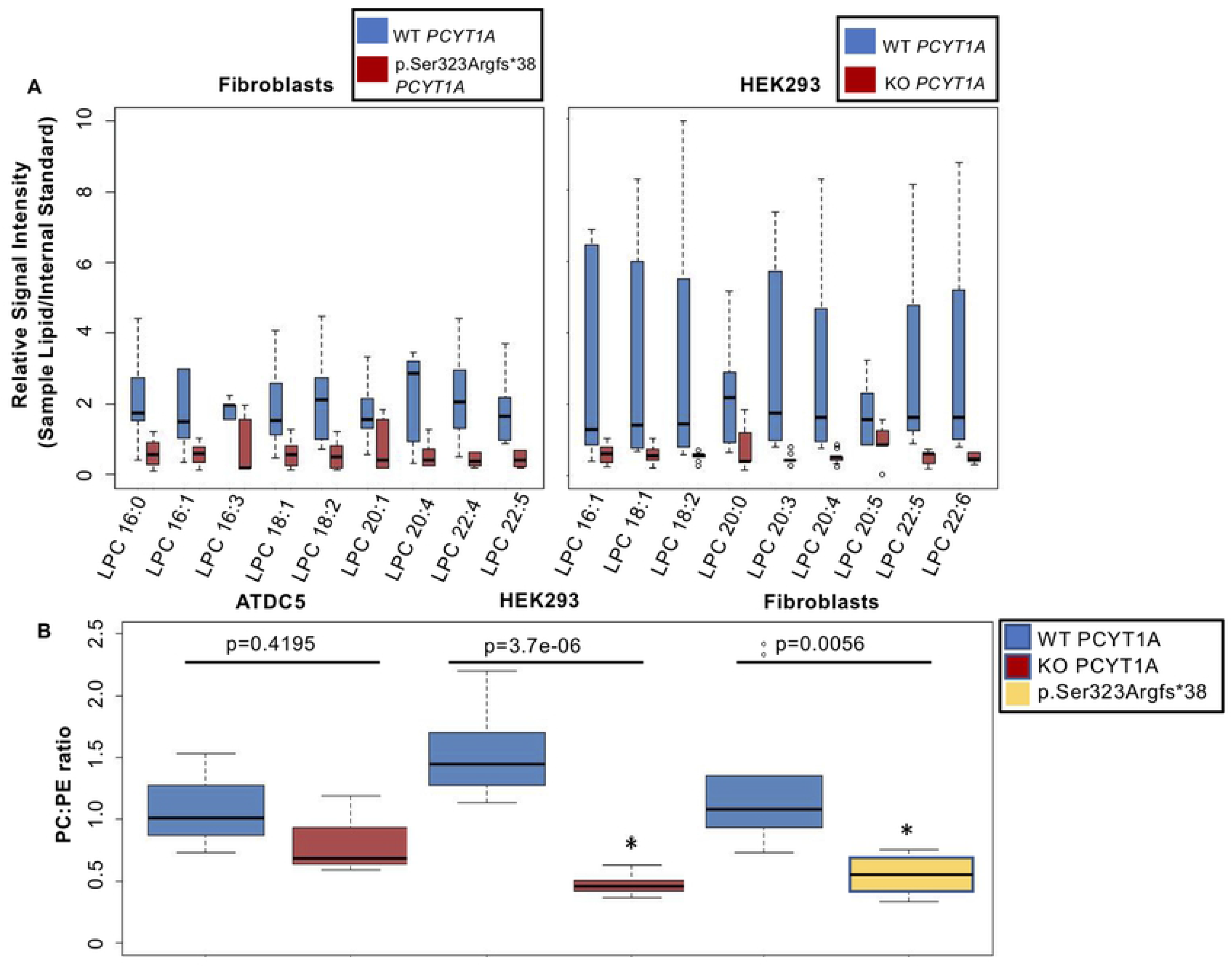
Lipidomics reveal decreases in LPCs and in PC:PE ratio in SMD-CRD patient fibroblasts and *PCYT1A*-null HEK293 cells. A) Data represent mean values of relative abundance of lysophosphatidylcholine (LPC) species of various acyl chain lengths and compositions relative to internal standards in SMD-CRD patient fibroblasts and *PCYT1A*-null HEK293 cells as compared to wild-type controls. Error bars represent SD of three replicates. p<0.05 for all comparisons. B) Ratios of total phosphatidylcholine (PC) to phosphatidylethanolamine (PE) in SMD-CRD patient fibroblasts, *PCYT1A*-null HEK293 cells, and *Pcyt1a*-null ATDC5 cells as compared to wild-type controls. Error bars represent SD. *p<0.05.

The levels of several LPC species (Fig 7A) were significantly decreased in *PCYT1A*-null HEK293 cells (LPCs 16:1, 18:1, 18:2, 20:0, 20:3, 20:4, 20:5, 22:5, and 22:6) and in SMD-CRD patient fibroblasts (LPCs 16:0, 16:1, 16:3, 18:1, 18:2, 20:1, 20:4, 22:4, and 22:5). Notably, levels of five identical LPCs (LPCs 16:1, 18:1, 18:2, 20:4, and 22:5) were reduced in both *PCYT1A*-null HEK293 cells and in SMD-CRD patient fibroblast cell lines as compared to controls (Fig 7A). The reduction over all LPC classes attributed to loss of *PCYT1A* or the p.Ser323Argfs*38 mutation was 3.3-fold for *PCYT1A*-null ATDC5 cells, 3.9-fold for *PCYT1A*-null HEK293 cells, and 1.7-fold for SMD-CRD fibroblasts. We also observed significant decreases in the lysophosphatidylethanolamines (1.5-fold in null ATDC5 cells) and increases in lysophosphatidylserines (2-fold in null HEK293 cells) (S2 Table, S4 Table).

PC:PE ratios were significantly reduced by 2-3 fold in *PCYT1A*-null HEK293 cells and in SMD-CRD patient fibroblast cell lines as compared to their wild-type counterparts (Fig 7B). The PC:PE ratio in *PCYT1A*-null ATDC5 cells was reduced, but the reduction was not statistically significant. Because PC is a precursor to sphingomyelin (SM), we probed for relative and absolute changes in the levels of SM and its metabolite, ceramide (Cer), which has been implicated in apoptotic processes [34]. We found that sphingomyelin:ceramide (SM:Cer) ratios were not significantly altered in any of the *PCYT1A*-null or SMD-CRD patient cell lines as compared to their wild-type counterparts (S3 Fig). Other interesting lipidomic changes included a 1.4-fold increase in diacylglycerol in SMD-CRD cells (S3 Table), in line with a reduced rate of PC synthesis. The *PCYT1A*-null HEK293 lipidome showed significantly increased galactosylceramide and lactosylceramide (1.3-fold) (S4 Table).

## Discussion

### *PCYT1A* expression and PC synthesis rates do not correlate well with phenotypic severity

Our results show that all SMD-CRD patient-derived dermal fibroblasts tested have reduced steady state CCTα levels, ranging from 10 to 75% of control steady state levels, and reduced PC synthesis, ranging from 22 to 54% of wild-type. Surprisingly, the extent of reduction in steady-state CCTα amount or PC synthesis does not correlate with the phenotypic severity. Patients homozygous for p.Ala99Val had a severe phenotype despite the mildest fibroblast biochemical phenotype of all variants tested, both in their CCTα steady state levels (75% of wild-type) and PC incorporation (54% of wild-type). Similar levels of reduction in PC synthesis were independently observed in a recent study of *PCYT1A*-null Caco2 cells [33], which suggested that residual PC production could be attributable to CCTβ. The PC synthesis assays monitored radiolabeled choline uptake into PC in a 2 hr pulse, but this value reflects the balance between incorporation and loss due to PC degradation. Lipidomic analyses revealed that CCTα deficiency in HEK293 cells can suppress PC degradation. Nonetheless, because the p.Ala99Val allele decreased PC incorporation more severely than steady-state expression, we can conclude that this allele reduces the catalytic activity of CCTα independently of its effects on protein levels. This result is supported by recent kinetic studies of purified CCTα, which demonstrate impaired catalytic activity of p.Ala99Val and other SMD-CRD alleles in the catalytic domain [25]. In the crystal structure of the orthologous rat CCTα protein, this alanine inserts into a hydrophobic pocket that cannot easily accommodate a bulkier residue such as valine and leads to lower thermal stability [25,35], so the consequences of this allele are also consistent with our expectations based on the structure of the enzyme.

Other alleles reduced steady state levels of CCTα and PC incorporation much more severely. For instance, fibroblasts homozygous for the p.Ser323Argfs*38 allele had only 10% of the steady state CCTα and 22% of the PC incorporation levels of wild-type fibroblasts. We did not expect this allele to result in complete loss of CCTα expression, since the generated termination codon is C-terminal to the penultimate exon junction and therefore is predicted to escape nonsense-mediated mRNA decay [36]. However, the reduced steady state level of CCTα with the reduction in PC incorporation suggests that this allele translates into an unstable protein, likely due to its foreign P-region amino acid sequence.

Fibroblasts harboring compound heterozygous p.Ser323Argfs*38/p.Ser114Thr alleles had only 14% of wild-type steady state CCTα but 33% of wild-type PC incorporation. The protein levels in this cell line were only slightly higher than those of p.Ser323Argfs*38 homozygotes, indicating that the p.Ser114Thr allele has similarly adverse consequences on protein production and/or stability, at least when in combination with the p.Ser323Argfs*38 allele. This is consistent with findings from Cornell et al. (2019), showing that cDNA transfection of p.Ser114Thr into COS cells generated no signal above background, unlike transfection with WT *PCYT1A* or other alleles, perhaps due to misfolding and subsequent degradation [25]. Finally, fibroblasts homozygous for p.Glu129Lys had ~30% of wild-type steady-state CCTα levels and similar levels of PC incorporation (22% of wild-type). This glutamic acid residue is pivotal in hydrogen-bonding interactions between two α-helices of the catalytic domain Rossman fold, and the p.Glu129Lys mutation destabilizes that domain, lowering the T_m_ for unfolding by 6°C [25,35].

### CCT-associated changes in LD morphology may require complete ablation of its activity

Contrary to our expectations based on CCTα gene silencing in *Drosophila*, mouse, rat, and human cell lines [24,27,32,33], SMD-CRD patient fibroblasts (p.Ser323Argfs*38 homozygotes) failed to show significant differences in lipid droplet numbers or sizes as compared to controls, indicating that oleate loading does not induce an abnormal lipid droplet phenotype in cells with this genotype. Of note, p.Ser323Argfs*38 homozygous fibroblasts had the most severe reductions of all SMD-CRD patient cells tested in both steady-state levels of CCTα (~10% wild-type levels) and in phosphatidylcholine incorporation (~22% wild-type levels), so the absence of a lipid droplet phenotype in our experiments is surprising. This may be due to the lack of conservation of the mechanism for regulating lipid droplet size and number in human fibroblasts as compared to the models used in previous studies, including *Drosophila* S2 cells and larval fat bodies; mouse bone marrow-derived mouse macrophages and 3T3-L1 cells (embryonic fibroblasts that differentiate into adipocyte-like cells); rat IEC-18 cells (epithelial ileum); and human Caco2 cells (epithelial colorectal adenocarcinoma) [24,27,32,33]. Alternatively, it is possible that complete ablation of CCTα protein expression is required for lipid droplet phenotypes to manifest. The SMD-CRD patient fibroblasts retained some residual CCTα expression and activity, which could be sufficient for normal lipid droplet formation. Consistent with the above-mentioned studies in other cell types and a requirement for total ablation of CCT expression, *PCYT1A*-null HEK293 cells generated fewer and larger lipid droplets than did wild-type HEK293 cells in response to oleate loading. Transfection with wild-type, p.Ala99Val, and p.Tyr240His *PCYT1A* cDNAs rescued lipid droplet phenotypes in *PCYT1A*-null cells. This suggests that these variants in the catalytic or membrane-binding domains of CCTα provide ample enzyme activity and PC production to generate lipid droplets normally; this interpretation is further supported by our findings in SMD-CRD patient fibroblasts, in which even small residual amounts of functional CCTα are sufficient to form normal lipid droplets. However, we also recognize that cDNA expression was driven by a CMV promoter in our transfection studies, which may simply normalize lipid droplet phenotypes by driving *PCYT1A* overexpression. Analysis of the impact of the *PCYT1A* alleles associated with lipodystrophy on the size and number of LDs in patient cells has not been done, but could help unravel the basis of the differential disease manifestations linked to specific alleles.

### *PCYT1A*-null cells have grossly normal morphology and proliferation rates

Given that homozygous *Pcyt1a*-null mice are embryonic lethal; CHO MT-58 cells undergo apoptosis; and yeast cells reliant on M-domain or catalytic domain mutant *pct1* for PC synthesis show reduced viability [9,11–13], we are intrigued that *PCYT1A*-null HEK293 and ATDC5 cells are grossly morphologically normal and proliferate at rates comparable to their wild-type counterparts. This is contrary to recent work demonstrating reduced proliferation of *PCYT1A*-null Caco2 cells [33]. Our results are consistent with findings that *Pcyt1a*-null mouse peritoneal macrophages grow normally under standard cell culture conditions due to compensatory upregulation of a specific isoform of *Pcyt1b* [37]. Interestingly, these *Pcyt1a*-null mouse macrophages have increased susceptibility to cell death upon loading with free cholesterol, suggesting that additional stressors may be required to evoke cell death in CCTα-deficient cell lines. Furthermore, we show that *PCYT1A*-null HEK293 cells proliferate more quickly than their wild-type counterparts when cultured in delipidized serum. Our results suggest that growth rates are normalized independently of exogenous lipid uptake from serum in these cells and that *PCYT1A*-null HEK293 cells may have a growth advantage over their wild-type counterparts when grown in conditions devoid of exogenous lipids, perhaps by upregulation of compensatory pathways for PC production.

### Deficiency in *PCYT1A* expression and/or function reduces PC content and modifies PC remodeling

Using mass spectrometry, we investigated changes in lipid species that are directly implicated in or apart from those directly implicated in the Kennedy pathway. We found a broad array of lipids to be altered in *PCYT1A*-null HEK293 or ATDC5 cells or SMD-CRD patient fibroblast cell lines as compared to controls. Altered lipids were often cell-type specific. Key findings include lipid modifications that were replicated in two of the three cell lines tested. First, total PC:PE ratios were significantly decreased in *PCYT1A*-null HEK293 and in SMD-CRD patient fibroblast cell lines as compared to controls. CCTα deficiency manifested as reduced PC:PE in several previous reports [3,13,38]. Decreased PC:PE ratios have previously been associated with loss of membrane integrity, decreases in membrane potential, and membrane packing defects due to increased proportions of conical lipids leading to increased membrane stress [13,38]. Our data suggest that decreased PC:PE ratios leads to adverse consequences in SMD-CRD patients, perhaps in combination with the other defects we describe here.

The only lipid whose overall cellular content was decreased in two cell types was LPC, which was present in lower proportions in *PCYT1A*-null HEK293 and ATDC5 cell lines as compared to corresponding wild-type controls. In *Pcyt1a*-null ATDC5 cells, we also observed decreases in overall LPE content, suggesting potential upregulation of LPE acyltransferase (LPEAT) or downregulation of PLA_2_. In *PCYT1A*-null HEK293 cells, we observed decreased PC content, suggesting that compensatory pathways for PC production may not be as robust in this cell type. *PCYT1A*-null HEK293 cells also had increased content of LPS, galactosylceramide, and lactosylceramide. The implications of increased levels of these lipids are unclear. Both galactosylceramide and lactosylceramide are formed from ceramide, which can combine with PC to form sphingomyelin in a reaction catalyzed by sphingomyelin synthase. It is possible that decreased PC production by the Kennedy pathway depresses PC conversion to sphingomyelin in an attempt to maintain cellular PC, which in turn shunts ceramide into galactosylceramide and lactosylceramide production. In homozygous p.Ser323Argfs*38 fibroblasts, the proportions of DAG and PE were increased as compared to wild-type fibroblasts. We expected increased proportions of DAG, given that SMD-CRD cells have decreased production of CDP-choline, which normally combines with DAG to form PC in the final step of the Kennedy pathway. However, increased proportions of PE were not expected *a priori.* The increase in PE content could result from shunting the DAG that is non-utilized for PC synthesis into the CDP-ethanolamine pathway. We suggest that in SMD-CRD fibroblasts, upregulation of the PEMT pathway is not a major means for compensatory PC production.

In addition, we found that several LPCs were significantly decreased in both *PCYT1A*-null HEK293 and SMD-CRD patient fibroblast cell lines as compared to the corresponding control cell lines. Decreased LPCs could be attributable either to increased conversion of LPC into PC by LPCAT or to decreased conversion of PC into LPC by PLA_2_ (Fig 1). The former mechanism is supported by recent work by Lee and Ridgway (2018), who showed rescue in lipid droplet phenotypes in *PCYT1A*-null Caco2 cells supplemented with LPC [33]. Previous studies in COS cells and in CHO cells have shown that CCTα overexpression increases LPC levels by promoting PLA_2_-mediated PC catabolism [39,40]. It is possible that decreased activity by this pathway is also responsible for decreases in LPC in our cell lines.

Interestingly, recent studies have also demonstrated increased *Lpcat4* mRNA expression and activity toward LPC acyl chains 18:1, 18:2, 20:4, and 22:6 during the late stages of chondrocyte differentiation in ATDC5 cells [41]. Notably, LPCs with three of these four acyl chains (18:1, 18:2, and 20:4) were decreased in SMD-CRD patient fibroblast cell lines, and all four were decreased in *PCYT1A*-null HEK293 cells as compared to wild-type controls. The mean levels of these and other individual LPCs were also decreased in *Pcyt1a*-null as compared to wild-type ATDC5 cells, but none of these reductions approached statistical significance. This may be due in part to the small sample size of ATDC5 cells measured in our study. Further investigation of LPCs in this cell type is indicated, but we posit that decreased LPCs may play a role in SMD-CRD pathophysiology.

Increased LPCAT expression or activity is also suggested by our finding that *Pcyt1a*-null ATDC5 cells have increased rates of chondrocyte differentiation as compared to wild-type controls. *Lpcat4* knockdown has been associated with decreased rates of chondrocyte differentiation in ATDC5 cells [41]. Compensatory upregulation of LPCAT proteins may contribute to the increased rate of chondrocyte differentiation in *Pcyt1a*-null ATDC5 cells. Changes in chondrocyte differentiation rates have also been observed in previous studies of other skeletal dysplasias. For example, ATDC5 cells expressing FGFR3-G380R, the variant responsible for achondroplasia, have reduced rates of chondrocyte differentiation, whereas ATDC5 cells harboring inactivating mutations in the Kabuki syndrome gene *Kmt2d* show precocious differentiation [42,43]. Our results suggest that SMD-CRD may be another phenotype for which perturbed rates of chondrocyte differentiation contribute to skeletal dysplasia.

Overall, our results indicate that partial or complete loss of *PCYT1A* in ATDC5, HEK293, or fibroblast cell lines leads to decreased PC synthesis by the Kennedy pathway and cell-type specific secondary dysregulation of diverse lipid metabolic pathways and phenotypes. Our results also suggest involvement of *Pcyt1a* in chondrocyte differentiation but not proliferation or morphology. We recommend a careful assessment of all *PCYT1A* alleles implicated in SMD-CRD, isolated retinopathy, and lipodystrophy in a range of cell types to dissect further tissue-specific consequences of these alleles.

## Materials and Methods

### Patient consent

Our study was approved by the Johns Hopkins Medicine Institutional Review Board and by the IRBs of other participating institutions. We obtained informed consent from all responsible individuals who participated in this study.

### Cell culture

Primary adherent fibroblast cells were derived from skin biopsies of SMD-CRD patients or wild-type controls. HEK293 cells were a gift from Dr. Jeremy Nathans of Johns Hopkins University. ATDC5 cells were a gift from Dr. Jill Fahrner of Johns Hopkins University (ECACC, Cat#99072806). All cell lines were cultured at 37° C under 5% CO_2_ and passaged using 1x trypsin-EDTA (Gibco, Cat#15400-054). Fibroblast, HEK293, and CHO cell lines were cultured in 1x MEM (Gibco, Cat#11430-030) supplemented with non-essential amino acids (Gibco, Cat#11140-050), 1% L-glutamine (Gibco, Cat#25030-149), and 10% fetal bovine serum (FBS; Gemini Biosciences, Cat#100-106). For experiments using delipidated serum, culture medium was the same as above, except 10% delipidated FBS (Gemini Biosciences, Cat#900-123) was substituted for regular FBS.

Prior to chondrocyte differentiation, ATDC5 cells were maintained in DMEM/F12 50/50 (Corning Cellgro, Cat#15-090-CV) with 1% L-glutamine (Gibco, Cat#25030-149), and 5% FBS (Gemini Biosciences, Cat#100-106), and 1% penicillin-streptomycin (Gibco, Cat#15140-122). Chondrocyte differentiation was induced by adding 1x insulin/transferrin/selenium (Corning Cellgro, Cat#25-800-CR) to the culture medium for 21 days.

### DNA extraction and Sanger sequencing

DNA was extracted from blood or cell lines using the Puregene Blood Core Kit B (Qiagen, Cat#158467). *PCYT1A* exons were PCR-amplified from SMD-CRD patient genomic DNA with Accuprime Taq polymerase (Invitrogen, Cat#12339-016) and standard thermal cycling conditions using custom primer sequences (S1 Table).

### *PCYT1A* cDNA construct generation

RNA was extracted from *PCYT1A*-wild-type human fibroblasts using the RNeasy Mini Kit (Qiagen, Cat#74104), reverse-transcribed according to the manufacturer’s protocol (SuperScriptIII One-Step RT-PCR system with Platinum Taq DNA Polymerase;Invitrogen, Cat# 12574-018, primer sequences in S1 Table), and cloned into the pcDNA3 mammalian expression vector (Invitrogen, Cat#V790-20) using primers designed to amplify full-length *PCYT1A* cDNA (based on NCBI transcript NM_005017.2) and attach restriction enzyme sites for subsequent cloning (forward primer: 5’-TACGTAAGCTTAGCGCCACCTCAGAAGATAA-3’, reverse primer: 5’-CTGAGCTCGAGTTGGGGTCACAATTTGGAAT-3’). Single base-pair SMD-CRD missense variants were then introduced using QuikChange II XL Site-Directed Mutagenesis Kit (Agilent, Cat # 200521; primer sequences in S1 Table). All plasmids were sequence-verified prior to use.

### Plasmid transfection

Transfections were conducted using Lipofectamine 2000 as prescribed by the manufacturer’s protocol (Invitrogen, Cat#11668-019) with ratios of 1.6 μg total cDNA-containing plasmid: 4 μL Lipofectamine for 12-well plate transfections, or ratios of 4.0 μg total plasmid DNA: 10 μL Lipofectamine for 6-well plate transfections. For lipid droplet rescue experiments, plasmid cDNAs harboring wild-type or SMD-CRD patient variants in *PCYT1A* were transfected into *PCYT1A*-null HEK293 cells and 24 hours post-transfection, cells were incubated with 1 mM oleate medium and further processed for imaging, as described below.

### Protein extraction and immunoblotting

At approximately 80% confluency, we washed monolayers with 1x PBS and scraped cells into ice-cold 1xPBS supplemented with protease inhibitors (Sigma-Aldrich, Cat#P8340-5ML), and centrifuged them for 5 minutes at 1,000 x g. We then resuspended the cells in RIPA buffer (150 mM NaCl, 1.0% NP-40, 0.5% sodium deoxycholate, 0.1% SDS, 50 mM Tris, pH 8.0) with protease inhibitors (Sigma-Aldrich, Cat#P8340-5ML), vortexed briefly, and lysed the cells at 4° C with rotation and periodic low-speed vortexing. We pelleted cell debris by centrifugation at 12,000 x g for 15 minutes and isolated the supernatant for subsequent analysis. We measured total protein concentrations of cell lysates with the BCA protein assay kit (Pierce, Cat#23225) and diluted 40 μg of each protein with XT sample buffer (Bio-Rad, Cat#1610791) and XT reducing agent (Bio-Rad, Cat#1610792) as prescribed by company protocols. Samples were denatured by boiling for 5 minutes, then cooled and run on 4-12% Bis-Tris Criterion XT precast gels (Bio-Rad, Cat#3450123) with XT MOPS Running Buffer (Bio-Rad, Cat#1610788). Transfer was conducted onto Immun-Blot PVDF membranes (Bio-Rad, Cat#162-0177) at 4° C with 100 V for 1 hour in 1 x Tris/Glycine Buffer (Bio-Rad, Cat#1610734) and 20% methanol (Fisher Scientific, Cat#A41220). Blots were blocked for 1 hour at room temperature in 5% nonfat dried milk in TBST (0.5% Tween-20, 137 mM NaCl, 200 mM Tris, pH 7.5), then incubated in the appropriate primary antibodies (anti-β-actin antibody AC-15, ThermoFisher Scientific, Cat#AM4302, 1:10,000 dilution; monoclonal anti-CCTα, Abcam, Cat#ab109263, 1:1,000 dilution) diluted in 5% nonfat dried milk in TBST at 4° C overnight. The epitope for this antibody is N-terminal to the SMD-CRD mutations, and thus should react equally with both WT and mutant CCTα species. Membranes were washed 3 x 10 minutes in TBST, incubated for 1 hour at room temperature in the appropriate secondary antibodies diluted 1:10,000 in 5% nonfat dried milk in TBST (goat anti-rabbit IgG-HRP, Santa Cruz Biotechnology, Cat#sc-2004; goat anti-mouse IgG-HRP, Santa Cruz Biotechnology, Cat#sc-2005). Membranes were washed 3 x 10 minutes in TBST, then incubated in ECL reagent (GE Healthcare Life Sciences, Cat#RPN2106) for 5 minutes and exposed to CLX Posure Film (Thermo Scientific, Cat#34091).

### 3H-choline incorporation assay

To assess the effect of *PCYT1A* variants on the rate of phosphatidylcholine synthesis by the Kennedy pathway, we monitored continuous incorporation of [Methyl-^3^H]-choline chloride (NEN Radiochemicals, Cat#NET109001MC) into phosphatidylcholine over time essentially as described previously [44]. We cultured control or SMD-CRD patient fibroblasts in 1x MEM with 10% FBS, 1% L-glutamine, and non-essential amino acids in 60 mm dishes until ~80% confluency and counted a subset of cells using the Beckman Coulter Particle Counter Z1. We aspirated the normal growth medium from remaining cells and incubated in serum-free medium at 37° C with 5% CO_2_ for 1 hour. We then added 5 µCi [^3^H]-methyl-choline per dish and returned to the incubator or quenched immediately for our 0-minute time point. Two hours later, we quickly rinsed cells in ice-cold 1xPBS and quenched radiolabel uptake by adding 1 mL of ice-cold methanol per dish. We then placed cells on ice and scraped into prechilled glass tubes containing 1 mL chloroform, and residual cells were scraped with an additional 1 mL of methanol. We then added 0.75 mL dI water per tube, mixed by vortexing at low speed, and added 1 mL additional chloroform followed by 1 mL of dI water per tube. We then vortexed cells briefly to mix, followed by centrifugation for 5 minutes at 1000 RPM to form a biphasic mixture. We transferred the lower organic phase (containing lipids) to a scintillation vial, evaporated under nitrogen gas (Organomation Associates, Inc., Berlin, MA, USA) and resuspended in 10 mL Insta-Gel Plus scintillation fluid (Perkin Elmer, Cat#6013399), followed by counting with a Beckman LS6500 scintillation counter. Previous thin layer chromatography analyses have shown that >90% of counts in the organic phase following up to 2 hours incubation are attributable to phosphatidylcholine [42], so we used counts in this phase as a proxy for radiolabeled phosphatidylcholine levels. We corrected all samples for background counts by subtracting counts generated at the 0 hour incubation timepoint from the 2 hour incubation timepoint.

### Genome editing

We performed genome editing as described previously, with slight modifications [46]. Guide RNAs against target genomic sequences from human or mouse *PCYT1A* (RefSeq NC_000003.11 or NC_000082.5, respectively) were designed using the CRISPR MIT portal [47] (S1 Table), modified into oligonucleotides for cloning purposes, and cloned according to the submitter’s laboratory protocol into the pSpCas9(BB)-2A-Puro (PX459) V2.0 plasmid (Addgene, Cat#62988), a gift from Feng Zhang’s laboratory [48]. By combining two guide RNAs targeting the beginning and end of the genomic sequence of either human or mouse *PCYT1A*, we generated ~31 kb genomic deletions in either human (HEK293 cells) or mouse (ATDC5 cells) *PCYT1A*, effectively excising the majority of the coding sequence. Cells were seeded for transfection at ~2 x 10^5^ cells/well in six-well plates, grown to ~80% confluency, and transfected using Lipofectamine 2000 as prescribed by the manufacturer’s protocol (Invitrogen, Cat#11668-019), with equal DNA amounts of each guide RNA-encoding plasmid (2 μg each plasmid DNA: 10 μl Lipofectamine). Next, we treated cells with puromycin-containing medium at either 2 μg/mL (HEK293 cells) or 4 μg/mL (ATDC5 cells) 24 hours post-transfection. Medium was replaced daily for 2-3 days, until untransfected control cells died. Single cells were then isolated by serial dilution in puromycin-free medium, expanded into isogenic clones, subjected to DNA extraction, and screened for both wild-type and deletion-containing (S1 Table) genomic DNA by PCR amplification. Clones which contained the *PCYT1A* deletion but not wild-type genomic DNA were genotyped using Sanger sequencing and tested for the presence of CCTα protein by immunoblot.

### Cell proliferation assays

We measured cell proliferation using the Cell Counting Kit-8 (Dojindo Molecular Technologies, Cat#CK04-05), as prescribed by the manufacturer’s protocol. We counted cells using the Beckman Coulter Particle Counter Z1, plated at 1000 cells per well in 96-well plates, and cultured for the respective amounts of time indicated in our growth curve. We measured growth by incubating with CCK-8 solution for 3 hours, followed by absorbance reading at 450 nm with the Biotek Synergy 2 plate reader and Gen5 software v2.01.14.

### Oleate incubation for lipid droplet and nuclear lamina translocation studies

In order to induce lipid droplet formation, we prepared oleic acid: BSA complexes for addition to cell culture medium. First, we sonicated 100 mM sodium oleate solution (Sigma-Aldrich, Cat#O3880-1G) in distilled water on ice (Branson Sonifier 250 with a tapered microtip, Fisher Scientific, Cat#22-309782) under a constant duty cycle and output level of 2 for approximately 2 minutes, or until solution reached clarity. We then mixed this solution with a 200 mg/ml fatty acid-free BSA (Sigma-Aldrich, Cat#A8806-5G) in PBS and briefly sonicated again until solution was clear. This generated a final stock of 100 mg/mL BSA with 10 mM oleate, which we diluted 10-fold in serum-free medium (1x MEM supplemented with non-essential amino acids and 1% L-glutamine) for a 1 mM working oleate solution. We incubated cells in 1 mM oleate medium for various timepoints prior to processing for immunofluorescence or lipid droplet staining.

### Cell staining and immunofluorescence

We performed cell culture using the conditions described above in 12-well plates containing sterile 18-mm glass cover slips (VWR, Cat#48380-046). We fixed cells for 10 minutes in 4% paraformaldehyde (Sigma-Aldrich, Cat#P6148-500G), permeabilized for 10 minutes with 0.2% Triton X-100 (Sigma-Aldrich, Cat#T-8532) in 1x PBS, and blocked for 1 hour in 1% BSA (Sigma-Aldrich, Cat#A-9647) in 1x PBS, all at room temperature. We then processed cells for immunofluorescence or lipid droplet staining.

For immunofluorescence experiments, we incubated cells in primary antibody (anti-CCTα, Abcam, Cat#ab109263) diluted 1:200 in 1% BSA in PBS overnight at 4°C, followed by a one-hour room-temperature incubation in Alexa Fluor 488 or 555 secondary antibodies (Thermo Fisher Scientific, Cat#A-21428 or Cat#A-11008) diluted 1:300 in 1% BSA in PBS. For lipid droplet staining, we incubated cells for one hour in BODIPY 493/503 dye (ThermoFisher Scientific, Cat#D-3922, stored as 1 mg/mL stock in 100% ethanol at −20°C) diluted in 1:500 in 1x PBS.

We counterstained cell nuclei with DAPI (Life Technologies, Cat#D1306) diluted to 300 nM in 1x PBS for 10 minutes at room temperature, followed by mounting with ProLong Gold Antifade Mountant (Thermo Fisher Scientific, Cat#P36930) on glass slides (Fisher Scientific, Cat#12-550-15). We obtained images using the Zeiss LSM 510 Meta Confocal Microscope under 63x oil magnification.

### Lipid droplet quantification

We quantified lipid droplet sizes and numbers as described previously [24]. Using ImageJ software (v1.47), we converted images to 8-bit, adjusted image threshold, and used the “analyze particle” command with a particle size distribution of 20 to infinity pixels^2 set to exclude on edges.

### Alcian blue staining

We performed Alcian blue staining as previously described [49], with slight modification. Upon reaching ~80% confluency, we rinsed cells in 1xPBS, fixed with 4% paraformaldehyde for 10 minutes at room temperature, and then incubated in 1% Alcian Blue 8GX (Sigma-Aldrich, Cat#A5268-10G) pH 1.0 at 4° C overnight. After briefly washing in 1xPBS, we lysed cells in 1% SDS for one hour and measured an absorbance of 605 nm using the Biotek Synergy 2 plate reader with Gen5 software v2.01.14.

### Lipid standards for mass spectrometry

We obtained the following lipid standards from Avanti Polar Lipids (Alabaster, AL): 1,2-dilauroyl-*sn*-glycero-3-phosphate (sodium salt) (PA 12:0/12:0, PA C12), 1,2-dilauroyl-*sn*-glycero-3-phosphocholine (PC 12:0/12:0, PC C12), 1,2-dilauroyl-*sn*-glycero-3-phosphoethanolamine (PE 12:0/12:0, PE C12), 1,2-dilauroyl-*sn*-glycero-3-phospho-(1’-*rac*-glycerol) (sodium salt) (PG 12:0/12:0, PG C12), 1,2-dilauroyl-*sn*-glycero-3-phospho-L-serine (sodium salt) (PS 12:0/12:0, PS C12), cholesteryl-d7 palmitate (Cholesterol-d7 ester 16:0, Cholesterol ester d7), N-lauroyl-D-*erythro*-sphingosine (C12 Ceramide d18:1/12:0, Cer C12), N-heptadecanoyl-D-erythro-sphingosine (C17 Ceramide d18:1/17:0, Cer C17), N-lauroyl-D-erythro-sphingosylphosphorylcholine (12:0 SM d18:1/12:0 SM C12), 1,3-dihexadecanoyl glycerol (d5) [1,3-16:0 DG (d5), DG d5], 1,3(d5)-dihexadecanoyl-2-octadecanoyl-glycerol [TG d5-(16:0/18:0/16:0), TG d5], D-galactosyl-ß-1,1’ N-lauroyl-D-*erythro*-sphingosine [C12 Galactosyl(ß) Ceramide (d18:1/12:0), GlcCer C12], D-lactosyl-ß-1,1’ N-dodecanoyl-D-*erythro*-sphingosine [Lactosyl (ß) C12 Ceramide, LacCer C12]. We purchased APCI positive calibration solution from AB Sciex (Cat. #4460131; Concord, Ontario, Canada). Lipid stocks were prepared in methanol, dichloromethane/methanol 1:1 (v/v), or dichloromethane and stored at −20°C. Ultrapure water (resistivity > 18 MΩ cm) was used for mass spectrometry experiments.

### Cell culture and lipid extraction for mass spectrometry

For mass spectrometry experiments, we measured various lipid classes in *PCYT1A* wild-type versus *PCYT1A*-null HEK293 and ATDC5 cells, or in cultured dermal fibroblasts obtained from healthy controls or SMD-CRD patients homozygous for the p.Ser323Argfs*38 variant in *PCYT1A*. All cell lines were cultured in growth conditions described above. Upon reaching approximately 80% confluency, cells were scraped in ultrapure water, pelleted at 1000 x g, resuspended in ultrapure water, and subjected to total protein concentration measurements using the BCA protein assay kit (Cat#23225, Pierce). We stored cell extracts at −80°C until further processing.

We obtained a total lipid extract from cell pellets using a modified Bligh-Dyer procedure [50]. In brief, we diluted each cell suspension (200 µg protein) to 200 µl with ddH_2_O and sonicated ten times using short bursts with a sonic homogenizer (Fisher Scientific, Waltham, MA) for 30 seconds followed by 30 seconds on ice. We transferred homogenates to glass tubes and gently mixed with 800 µL of ddH_2_O, followed by 2.9 mL methanol/dichloromethane (2:0.9, v/v) containing internal standards for 12 lipid classes to each sample. We then added 1 mL of ddH_2_O and 0.9 mL dichloromethane to obtain a biphasic mixture, which was incubated at 4°C for 30 min and centrifuged at 4°C for 10 min at 3000 x g to separate organic and aqueous phases. 1 mL of the organic phase was transferred to a 2 mL glass vial and stored at −20°C until use. We finally dried 0.5 mL of the organic layer extract using a nitrogen evaporator (Organomation Associates, Inc., Berlin, MA, USA), and re-suspended in 135 µL of running solvent (dichloromethane:methanol [1:1 v/v] with 5 mM ammonium acetate) containing 5 mg/mL of ceramide (C17:0) as an internal standard for instrument performance. All solvents used were HPLC grade.

### Lipidomic analysis by MS/MS^ALL^

In order to capture cellular changes in a broad spectrum of lipids, we performed mass spectrometry analyses on a TripleTOF™ 5600 mass spectrometer (AB SCIEX, Redwood City, CA) using an MS/MS^ALL^ approach. We introduced 50 µL of each sample extract into a DuoSpray electrospray ionization source at a flow rate of 7 µL/min. All samples were run in duplicate in positive ion mode. We used a mass resolution of ∼30,000 for TOF MS scans and ∼15,000 for product ion scans in high sensitivity mode, and the instrument automatically calibrated using an APCI positive calibration solution (AB Sciex) after every 10 sample injections. The source parameters were as follows: ion source gases 15 psi (GSI), 20 psi (GS2), curtain gas 30 psi, temperature 150°C, positive ion spray voltage +5200 V, declustering potential of 80V, and collision energy of 10V. Initial TOF MS scanning provided an overview of total lipid content at an accumulation time of 5s. Precursor ions were selected by sequential 1 Da mass steps from 200.050 to 1200.050 m/z, and analytes in each 1 Da step were introduced into the collision chamber. Fragments were identified by TOF with a scan range of 100-1500 m/z (accumulation time of 300 ms). The collision energy for each MS/MS step was 40 eV. Data were acquired using Analyst 1.7 TF (AB SCIEX, Concord, ON, Canada).

### Mass spectrometry data processing and analysis

Lipid identifications were pre-validated using a pooled sample that was extracted and sequentially run 8 times. Criteria for the inclusion of lipid analytes for analysis was that MS/MS fragment peaks were present in 7 of the 8 pooled runs, with a coefficient of variation (CV) for peak identifications less than 20%. Peak identifications meeting these criteria were then used to develop a targeted method in LipidView. The targeted method was used to identify these pre-validated lipid species in experimental samples using a custom made MatLab script and MultiQuant software (version 3.0, AB SCIEX, Concord, ON, Canada). All peak intensities were corrected by their corresponding internal standard, and each sample duplicate was averaged. If duplicates varied more than 30%, the sample was re-run. For statistical analyses, intensity values of 0 were replaced with a minimum intensity value that was calculated by dividing the average intensity value of for that particular lipid by 0.001. Data were log-transformed, z-score normalized, and exponentially transformed so that all values were positive. In order to compare each lipid between *PCYT1A* wild-type and variant cell lines of each group (ATDC5, HEK293, or fibroblast cell lines), we used Welch’s two-sample *t*-test.

### Statistical procedures

All statistical analyses were performed using R 3.2.4 (Bioconductor) [51]. Between-group comparisons were performed using Welch’s two-sample *t*-test. Multiple group comparisons were done using Tukey’s Honest Significant Difference test. No outliers were removed for statistical analyses.

## Acknowledgements

We are grateful to our patients and their families for participation in this study. We would like to thank Dr. Jill Fahrner of Johns Hopkins University for providing her ATDC5 cells.

## Supporting Information

**S1 Fig. Oleate induces CCTα membrane translocation in HEK293 cells, but not in control human fibroblasts. A)** HEK293 and **B)** CHO cells with and without oleate stimulation. LMNA-nuclear lamina marker. DAPI-nuclear marker.

**S2 Fig. Generation of** *PCYT1A***-null HEK293 and ATDC5 cells.** *PCYT1A*-null cell lines were generated by cotransfecting with guide RNAs positioned at the beginning and end of *PCYT1A* to introduce a 31-kb deletion that excises the majority of the coding sequence and ablates CCTα expression. **A)** Exon structure of *PCYT1A* (blue) with HEK293-targeting guide RNA placement (orange arrows) in the first and last coding exons. Tan arrows labeled A, B, and C indicate placement of forward and reverse primers used to sequence for WT or deleted alleles. A similar targeting strategy was used for editing in ATDC5 cells. **B)** Chromatogram of Cas9-induced deletion breakpoint in a genome-edited HEK293 cell clone. Both alleles contain the deletion, with one allele including an additional single-base insertion. **C-D)** Western blots showing ablation of endogenous CCTα expression in three separate clones of HEK293 cells **(C)** or ATDC5 cells **(D)** homozygous for a *PCYT1A* deletion.

**S3 Fig. SM:Cer ratios in WT and** *PCYT1A* **mutant cells.** Ratios of sphingomyelin (SM) to its metabolite ceramide (Cer) were not significantly altered in WT as compared to *PCYT1A*-null HEK293 and ATDC5 cell lines or in control as compared to homozygous *PCYT1A* p.S323Rfs*38 patient-derived cell lines. Data are represented as a box and whisker plot for which boxes represent the range of the first to third quartiles, lines represents median values, and whiskers extend to minimum and maximum data points.

**S1 Table. Oligo sequences for CRISPR experiments, site-directed mutagenesis, and sequencing of protein-coding exons of *PCYT1A.***

**S2 Table. Lipidomic analysis of WT or** *Pcyt1a***-null ATDC5 cells by MS/MS^ALL^**.

**S3 Table. Lipidomic analysis of WT or SMD-CRD patient-derived fibroblasts homozygous for the p.Ser323Argfs*38** *PCYT1A* **variant by MS/MS^ALL^**.

**S4 Table. Lipidomic analysis of WT or** *PCYT1A***-null HEK293 cells by MS/MS^ALL^.**

## References

1. Hoover-Fong J, Sobreira N, Jurgens J, Modaff P, Blout C, Moser A, et al. Mutations in PCYT1A, encoding a key regulator of phosphatidylcholine metabolism, cause spondylometaphyseal dysplasia with cone-rod dystrophy. Am J Hum Genetics. 2014;94:105–12.

2. Yamamoto GL, Baratela WA, Almeida TF, Lazar M, Afonso CL, Oyamada MK, et al. Mutations in PCYT1A cause spondylometaphyseal dysplasia with cone-rod dystrophy. Am J Hum Genetics. 2014;94:113–9.

3. Payne F, Lim K, Girousse A, Brown RJ, Kory N, Robbins A, et al. Mutations disrupting the Kennedy phosphatidylcholine pathway in humans with congenital lipodystrophy and fatty liver disease. Proc National Acad Sci. 2014;111:8901–6.

4. Testa F, Filippelli M, Brunetti-Pierri R, Fruscio G, Iorio V, Pizzo M, et al. Mutations in the PCYT1A gene are responsible for isolated forms of retinal dystrophy. Eur J Hum Genet. 2017;25:651–5.

5. Kennedy EP, Weiss SB. The function of cytidine coenzymes in the biosynthesis of phospholipids. Journal of Biological Chemistry. 1956;222:193–214.

6. Cornell RB, Ridgway ND. CTP:phosphocholine cytidylyltransferase: Function, regulation, and structure of an amphitropic enzyme required for membrane biogenesis. Progress in Lipid Research. 2015;59:147–71.

7. Karim M, Jackson P, Jackowski S. Gene structure, expression and identification of a new CTP:phosphocholine cytidylyltransferase β isoform. Biochimica et Biophysica Acta (BBA) - Molecular and Cell Biology of Lipids. 2003;1633:1–12.

8. Keenan TW, Morre JD. Phospholipid class and fatty acid composition of Golgi apparatus isolated from rat liver and comparison with other cell fractions. Biochemistry. 1970;9:19–25.

9. Wang L, Magdaleno S, Tabas I, Jackowski S. Early embryonic lethality in mice with targeted deletion of the CTP:phosphocholine cytidylyltransferase alpha gene (Pcyt1a). Mol Cell Biol. 2005;25:3357–63.

10. Carithers LJ, Ardlie K, Barcus M, Branton PA, Britton A, Buia SA, et al. A Novel Approach to High-Quality Postmortem Tissue Procurement: The GTEx Project. Biopreserv Biobank. 2015;13:311–9. V7 release, dbGaP Accession phs000424.v7.p2, accession date 12/04/17. https://www.gtexportal.org/.

11. Cui Z, Houweling M, Chen MH, Record M, Chap H, Vance DE, et al. A Genetic Defect in Phosphatidylcholine Biosynthesis Triggers Apoptosis in Chinese Hamster Ovary Cells. J Biol Chem. 1996;271:14668–71.

12. Sweitzer TD, Kent C. Expression of Wild-Type and Mutant Rat Liver CTP: Phosphocholine Cytidylyltransferase in a Cytidylyltransferase-Deficient Chinese Hamster Ovary Cell Line. Archives of Biochemistry and Biophysics. 1994;311:107–16.

13. Haider A, Wei Y-C, Lim K, Barbosa AD, Liu C-H, Weber U, et al. PCYT1A Regulates Phosphatidylcholine Homeostasis from the Inner Nuclear Membrane in Response to Membrane Stored Curvature Elastic Stress. Dev Cell. 2018;45:481–495.e8.

14. Jackowski S, Rehg JE, Zhang Y-M, Wang J, Miller K, Jackson P, et al. Disruption of CCTbeta2 expression leads to gonadal dysfunction. Mol Cell Biol. 2004;24:4720–33.

15. Hörl G, Wagner A, Cole LK, Malli R, Reicher H, Kotzbeck P, et al. Sequential synthesis and methylation of phosphatidylethanolamine promote lipid droplet biosynthesis and stability in tissue culture and in vivo. J Biol Chem. 2011;286:17338–50.

16. Lands WE. Metabolism of glycerolipides: A comparison of lecithin and triglyceride synthesis. Journal of Biological Chemistry. 1958;231:883–8.

17. Vance JE, Vance DE. Phospholipid biosynthesis in mammalian cells. Biochem Cell Biol. 2004;82:113–28.

18. Friedman JS, Chang B, Krauth DS, Lopez I, Waseem NH, Hurd RE, et al. Loss of lysophosphatidylcholine acyltransferase 1 leads to photoreceptor degeneration in rd11 mice. Proc National Acad Sci. 2010;107:15523–8.

19. Mitsuhashi S, Ohkuma A, Talim B, Karahashi M, Koumura T, Aoyama C, et al. A congenital muscular dystrophy with mitochondrial structural abnormalities caused by defective de novo phosphatidylcholine biosynthesis. Am J Hum Genetics. 2011;88:845–51.

20. Ahmed MY, Al-Khayat A, Al-Murshedi F, Al-Futaisi A, Chioza BA, Fernandez-Murray PJ, et al. A mutation of EPT1 (SELENOI) underlies a new disorder of Kennedy pathway phospholipid biosynthesis. Brain. 2017;140:547–54.

21. Vaz FM, rmott JH, Alders M, Wortmann SB, Kölker S, Pras-Raves ML, et al. Mutations in PCYT2 disrupt etherlipid biosynthesis and cause a complex hereditary spastic paraplegia. Brain. 2019;142:3382–97.

22. Dennis MK, Taneva SG, Cornell RB. The intrinsically disordered nuclear localization signal and phosphorylation segments distinguish the membrane affinity of two cytidylyltransferase isoforms. J Biol Chem. 2011;286:12349–60.

23. Gehrig K, Cornell RB, Ridgway ND. Expansion of the nucleoplasmic reticulum requires the coordinated activity of lamins and CTP:phosphocholine cytidylyltransferase alpha. Mol Biol Cell. 2008;19:237–47.

24. Aitchison AJ, Arsenault DJ, Ridgway ND. Nuclear-localized CTP:phosphocholine cytidylyltransferase α regulates phosphatidylcholine synthesis required for lipid droplet biogenesis. Mol Biol Cell. 2015;26:2927–38.

25. Cornell RB, Taneva SG, Dennis MK, Tse R, Dhillon RK, Lee J. Disease-linked mutations in the phosphatidylcholine regulatory enzyme CCTα impair enzymatic activity and fold stability. J Biol Chem. 2019;294:1490–501.

26. Walther TC, Chung J, Farese RV. Lipid Droplet Biogenesis. Annu Rev Cell Dev Bi. 2017;33:491–510.

27. Krahmer N, Guo Y, Wilfling F, Hilger M, Lingrell S, Heger K, et al. Phosphatidylcholine synthesis for lipid droplet expansion is mediated by localized activation of CTP:phosphocholine cytidylyltransferase. Cell Metab. 2011;14:504–15.

28. Cornell R, Vance DE. Translocation of CTP:phosphocholine cytidylyltransferase from cytosol to membranes in HeLa cells: stimulation by fatty acid, fatty alcohol, mono- and diacylglycerol. Biochimica et Biophysica Acta (BBA) - Lipids and Lipid Metabolism. 1987;919:26–36.

29. Lagace TA, Ridgway ND. The rate-limiting enzyme in phosphatidylcholine synthesis regulates proliferation of the nucleoplasmic reticulum. Mol Biol Cell. 2005;16:1120–30.

30. Pelech S, Pritchard P, Brindley D, Vance D. Fatty acids promote translocation of CTP:phosphocholine cytidylyltransferase to the endoplasmic reticulum and stimulate rat hepatic phosphatidylcholine synthesis. Journal of Biological Chemistry. 1983;258:6782–8.

31. Wang Y, MacDonald J, Kent C. Regulation of CTP:phosphocholine cytidylyltransferase in HeLa cells. Effect of oleate on phosphorylation and intracellular localization. Journal of Biological Chemistry. 1993;268:5512–8.

32. Guo Y, Walther TC, Rao M, Stuurman N, Goshima G, Terayama K, et al. Functional genomic screen reveals genes involved in lipid-droplet formation and utilization. Nature. 2008;453:657–61.

33. Lee J, Ridgway ND. Phosphatidylcholine synthesis regulates triglyceride storage and chylomicron secretion by Caco2 cells. J Lipid Res. 2018;59:1940–50.

34. Lozano J, Menendez S, Morales A, Ehleiter D, Liao W-C, Wagman R, et al. Cell Autonomous Apoptosis Defects in Acid Sphingomyelinase Knockout Fibroblasts. J Biol Chem. 2001;276:442–8.

35. Lee J, Johnson J, Ding Z, Paetzel M, Cornell RB. Crystal structure of a mammalian CTP: phosphocholine cytidylyltransferase catalytic domain reveals novel active site residues within a highly conserved nucleotidyltransferase fold. J Biol Chem. 2009;284:33535–48.

36. Popp M, Maquat LE. Organizing principles of mammalian nonsense-mediated mRNA decay. Annu Rev Genet. 2013;47:139–65.

37. Zhang D, Tang W, Yao P, Yang C, Xie B, Jackowski S, et al. Macrophages Deficient in CTP:Phosphocholine Cytidylyltransferase-α Are Viable under Normal Culture Conditions but Are Highly Susceptible to Free Cholesterol-induced Death: Molecular Genetic Evidence that the Induction of Phosphatidylcholine Biosynthesis in Free Cholesterol-Loaded Macrophages is an Adaptive Response. J Biol Chem. 2000;275:35368–76.

38. Li Z, Agellon LB, Allen TM, Umeda M, Jewell L, Mason A, et al. The ratio of phosphatidylcholine to phosphatidylethanolamine influences membrane integrity and steatohepatitis. Cell Metab. 2006;3:321–31.

39. Walkey C, Kalmar G, Cornell R. Overexpression of rat liver CTP:phosphocholine cytidylyltransferase accelerates phosphatidylcholine synthesis and degradation. Journal of Biological Chemistry. 1994;269:5742–9.

40. Barbour SE, Kapur A, Deal CL. Regulation of phosphatidylcholine homeostasis by calcium-independent phospholipase A2. Biochimica et Biophysica Acta (BBA) - Molecular and Cell Biology of Lipids. 1999;1439:77–88.

41. Tabe S, Hikiji H, Ariyoshi W, Hashidate-Yoshida T, Shindou H, Shimizu T, et al. Lysophosphatidylcholine acyltransferase 4 is involved in chondrogenic differentiation of ATDC5 cells. Sci Rep-uk. 2017;7:16701–16701.

42. Matsushita M, Kitoh H, Ohkawara B, Mishima K, Kaneko H, Ito M, et al. Meclozine facilitates proliferation and differentiation of chondrocytes by attenuating abnormally activated FGFR3 signaling in achondroplasia. Plos One. 2013;8:e81569–e81569.

43. Fahrner JA, Lin W-Y, Riddle RC, Boukas L, DeLeon VB, Chopra S, et al. Precocious chondrocyte differentiation disrupts skeletal growth in Kabuki syndrome mice. Jci Insight. 2019;4:e129380.

44. Kitos TE, Drobnies A, Ng M, Wen Y, Cornell RB. Contribution of lipid mediators to the regulation of phosphatidylcholine synthesis by angiotensin. Biochimica et Biophysica Acta (BBA) - Molecular and Cell Biology of Lipids. 2006;1761:261–71.

45. Ng M, Kitos TE, Cornell RB. Contribution of lipid second messengers to the regulation of phosphatidylcholine synthesis during cell cycle re-entry. Biochimica et Biophysica Acta (BBA) - Molecular and Cell Biology of Lipids. 2004;1686:85–99.

46. Moyer TC, Holland AJ. Generation of a conditional analog-sensitive kinase in human cells using CRISPR/Cas9-mediated genome engineering. Methods Cell Biol. 2015;129:19–36.

47. Hsu PD, Scott DA, Weinstein JA, Ran AF, Konermann S, Agarwala V, et al. DNA targeting specificity of RNA-guided Cas9 nucleases. Nat Biotechnol. 2013;31:827–32.

48. Ran AF, Hsu PD, Wright J, Agarwala V, Scott DA, Zhang F. Genome engineering using the CRISPR-Cas9 system. Nat Protoc. 2013;8:2281–308.

49. Lu C, Jain SU, Hoelper D, Bechet D, Molden RC, Ran L, et al. Histone H3K36 mutations promote sarcomagenesis through altered histone methylation landscape. Science. 2016;352:844–9.

50. Bligh E, Dyer W. A rapid method of total lipid extraction and purification. Can J Biochem Phys. 1959;37:911–7.

51. R Core Team. R: A Language and Environment for Statistical Computing. R Foundation for Statistical Computing, Vienna, Austria. 2018; https://www.R-project.org/.

